# Bigtools: a high-performance BigWig and BigBed library in Rust

**DOI:** 10.1101/2024.02.06.579187

**Authors:** Jack D. Huey, Nezar Abdennur

**Affiliations:** Program in Molecular Medicine, UMass Chan Medical School, Worcester, MA, 01605, USA; Department of Genomics and Computational Biology, UMass Chan Medical School, Worcester, MA, 01605, USA; Department of Systems Biology, UMass Chan Medical School, Worcester, MA, 01605, USA

## Abstract

The BigWig and BigBed file formats were originally designed for the visualization of next-generation sequencing data through a genome browser. Due to their versatility, these formats have long since become ubiquitous for the storage of processed sequencing data and regularly serve as the basis for downstream data analysis. As the number and size of sequencing experiments continues to accelerate, there is an increasing demand to efficiently generate and query BigWig and BigBed files in a scalable and robust manner, and to efficiently integrate these functionalities into data analysis environments and third-party applications. Here, we present *Bigtools*, a feature-complete, high-performance, and integrable software library for generating and querying both BigWig and BigBed files. *Bigtools* is written in the Rust programming language and includes a flexible suite of command line tools as well as bindings to Python. *Bigtools* is cross-platform and released under the MIT license. It is distributed on Crates.io and the Python Package Index, and the source code is available at https://github.com/jackh726/bigtools.

## Introduction

BigWig and BigBed, collectively referred to as Big Binary Indexed (BBI) files, are compressed, binary, and indexed formats for storing reference genome-aligned next-generation sequencing (NGS) information (1). BigBed is the binary equivalent of the Browser Extensible Data (BED) text format that represents features associated with (potentially overlapping) genomic intervals, e.g. peak calls or transcript annotations. Analogously, BigWig is the binary reincarnation of the textual Wiggle and BedGraph formats, which encode a quantitative step-function mapping unique floating point values to bases in the genome, while accommodating missing values, e.g. read pileups from a NGS experiment. Introduced in 2009, these file types are among the most common data artifacts derived from bulk and pseudo-bulk omic experimental modalities.

BBI files were designed for the display of large, distributed datasets in the UCSC Genome Browser; however, these features have made BigWig and BigBed useful beyond genome browsers, and today they are widely used for data analysis. For example, as of the end of 2023, the ENCODE Project data portal (2) provides for analysis purposes approximately 15,000 BigWig files and 17,000 BigBed files comprising 17 different schemas, excluding archived data. In BBI files, the primary data, index, and summary data at several resolutions or “zoom levels” are stored within the same file. The block compression and sparse index design allows for efficient random and remote access. However, because of their complex binary architecture, BBI files require specialized supporting software to enable reading, writing, and random access.

As the availability and use of BBI files grows, there is a growing desire to integrate BBI functionality into thirdparty bioinformatic applications. Since their inception, the command line interface (CLI) binaries from Kent et al. (1), hereon referred to as the UCSC tools, have served as the canonical software for generating and interacting with BigWig and BigBed files. While it is possible to interface directly with the C code that powers the UCSC tool executables (e.g. *bx-python* (3), *rtracklayer* (4), *bwtool* (5), *plastid* (6), and *pybbi* (7)), architectural limitations hinder its suitability as a standalone BBI library. This prevents widespread use of the UCSC reference implementation for application programming or for integration in interactive computing environments, particularly for languages with a runtime like Python, R, or Javascript.

Aside from the need for third-party integration, there is also a growing need for increased computational flexibility to tailor to different environments and use cases. First, while BBI files have traditionally been served statically on web servers, today users commonly require fast and easy querying of data locally for bioinformatic analyses. Similarly, data processing and generation of BBI files must also scale with rising volumes of raw NGS data. While traditionally done on highperformance computing clusters, data processing is also expanding into cloud compute platforms, particularly in large research consortia. In such environments, optimizing compute resources directly translates to decreased operating costs as well as reduced environmental impact.

We introduce *Bigtools*, a full-featured Rust library for BBI file creation, access, and manipulation that supports flexible resource usage and parallelization. Leveraging this library, *Bigtools* also includes versatile and high-performance CLI tools and Python bindings.

## Results

### Bigtools is a full-featured BBI library

Due to the limitations of the UCSC reference implementation, a number of alternative BBI software implementations have emerged in different languages, having varying levels of feature support and caveats (*libBigWig* and *pyBigWig* (8, 9), *bbi*.*js* (10), *Big* (11), *bigwig-reader* (12), see Table 1). These implementations are most commonly limited by either a lack of support for BigBed files (reading or writing), or both a lack of support for writing BBI files altogether. The library with the most complete support for reading and writing both file formats, *Big*, is written in Kotlin and thus restricted to the Java Virtual Machine (JVM) runtime and JVM-based languages. *Bigtools* is an independent and full-featured library for efficient creation, access, and manipulation of BBI files. It provides complete read and write support for both BigWig and BigBed files. *Bigtools* supports (i) granular random access to BBI files, including access to BBI file metadata, zoom levels, and summary records, (ii) interpreting custom BigBed records using embedded schemas written in the *autoSql* definition language (13), (iii) parallelization of reading and writing over multiple threads or cores, and (iv) optional memory-efficient single-pass creation of BBI files. The latter feature eliminates the requirements to start from a text representation, allowing for writing from standard input or progressive generation of data programmatically.

**Table 1.**
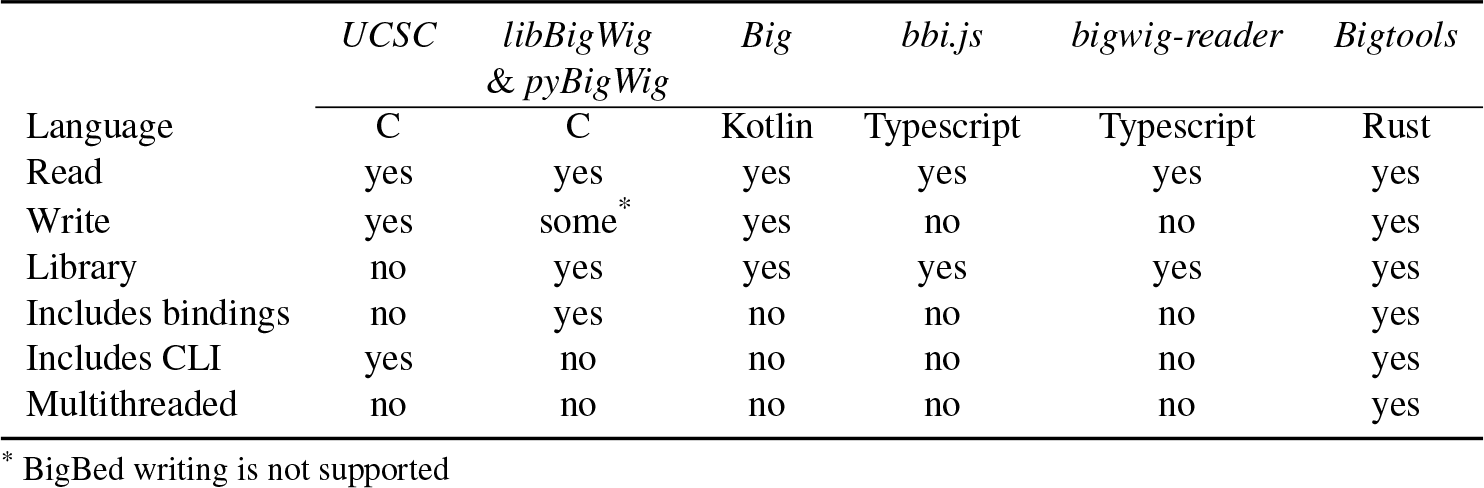
Comparison of Big Binary Indexed file software implementations.

*Bigtools* is written in Rust, which offers modern packaging and development tooling alongside its guarantees of memory-safety. Importantly, *Bigtools* can be effortlessly used by other Rust libraries and binaries through the Cargo package manager and `crates.io` package registry (https://doc.rust-lang.org/cargo/). Additionally, because Rust allows creation of C-compatible data structures and functions that can be called through any language that offers C Application Binary Interface (ABI) compatibility, there exist many Rust packages that make it easy to create and distribute bindings for commonly used languages. Along with the *Bigtools* library and CLI tools, we provide a companion Python library, *Pybigtools*, built using the *PyO3* crate (https://pyo3.rs/). Similar to the *pyBigWig* bindings for the *libBigWig* C library (9), this Python library exposes the I/O features of the *Bigtools* Rust library to Python and interfaces directly with efficient data structures in Python, including *NumPy* (14) arrays.

### Bigtools offers high-performance and flexible CLI tools

While alternative implementations for reading and writing BigWig and BigBed files are popular, none so far offer alternative CLI tools. This limits their use, particularly in pipelines and scripts. Here, we show that the *Bigtools* CLI has not only substantial performance improvements over the reference implementation but also a number of ergonomic benefits that can make it more versatile and easier to use.

To measure differences in performance between *Bigtools* and the reference implementation, we have set up a number of benchmarks (Table 2) comparing the processing of real-world data available from the ENCODE consortium (15). Each benchmark primarily measures the run time and maximum memory, but also samples CPU usage and memory over time. The benchmarks are repeated three times.

**Table 2.** Performance comparison between bigtools and UCSC tools.

| Benchmark | Wall time (sec) |  |  | Max memory (MB) |  |  |
| --- | --- | --- | --- | --- | --- | --- |
|  | UCSC | Bigtools ST | Bigtools MT | UCSC | Bigtools ST | Bigtools MT |
| bigwigaverageoverbed | 15.2±0.5 | 6.2±0.1 | 2.7±0.0 | 2083.5±0.1 | 74.8±0.1 | 233.8±10.7 |
| bigwigmerge | 113.1±1.6 | 65.5±1.8 | N/A | 2975.3±0.1 | 8.7±0.1 | N/A |
| bedgraphtobigwig | 93.8±1.5 | 47.2±1.1 | 16.4±0.2 | 218.8±0.1 | 23.5±0.1 | 50.6±1.0 |
| bedtobigbed | 61.7±0.2 | 37.5±1.5 | 28.8±0.1 | 366.7±0.2 | 52.5±0.0 | 32.8±0.3 |

ST: Single-threaded; MT: Multi-threaded (6 threads)

First, we compare single-threaded performance between *Bigtools* and the UCSC implementation. The wall time of *Bigtools* varies between approximately 1.5*×* to 2.5*×* faster than the UCSC tools. In addition, the *Bigtools* tools use considerably less memory (approximately 7*×* to 340*×*) than the UCSC tools. Notably, for tools like `bigWigAverageOverBed` and `bigWigMerge`, where the UCSC implementation loads in whole chromosomes at a time, *Bigtools* implements lazy computation which reduces the memory consumption by over an order of magnitude.

Second, unlike the UCSC tools, *Bigtools* offers multithreaded implementations for nearly all CLI tools (excluding only `bigwigmerge`). Increasing the number of threads to just two results in between approximately 3% to 41% faster wall time, while scaling up to six threads results in between approximately 24% to 73% faster wall time (Table 2). Memory usage does increase moderately for most of the tools, but remains comfortably below that of the UCSC tools.

Aside from performance, *Bigtools* offers a number of ergonomic benefits over the UCSC implementation. First, *Bigtools* offers several options for configuration that allow flexibility over the input representation. For example, while writing a BigWig file, the UCSC implementation reads through the input bedGraph file multiple times, which prohibits creating a BigWig from standard input. By contrast, *Bigtools* provides the option to write BBI files in a single pass of the input data, including from standard input. These configuration options also allow tuning of performance to the size of the input data: multiple passes are fine for smaller files, but results in significant time overhead for very large files. While writing as a single-pass *Bigtools* utilizes temporary files to avoid a memory overhead from index and zoom building, though this can be disabled. In addition to various configuration options, *Bigtools* also works as a drop-in replacement for the equivalent UCSC tools, supporting the most commonly used CLI flags. Finally, *Bigtools* can be used either as a single binary with each tool as a subcommand or as separate binaries for each tool.

## Conclusion

In summary, we present here a full-featured and highperformance library written in Rust for reading and writing BigWig and BigBed files, along with associated CLI tools and a Python library. *Bigtools* adds to the growing arsenal of core NGS tools in the Rust ecosystem (16, 17). It offers a number of configuration options that allow flexibility depending on file size and resource requirements, including scaling to multiple cores. Thus, *Bigtools* is ideal not only for local use where users may want to prioritize quick, parallel processing in return for higher processor and memory use, but also for cloud or batch workflows where users may want to prioritize minimal system resources in return for slightly longer computation.

*Bigtools* offers substantial benefits over previous library implementations of BigWig and BigBed reading/writing. Namely, *Bigtools* provides: (i) all features available in the UCSC tools, (ii) ease of integration and cross-language compatibility, and (iii) improved performance. In fact, during its development, it has already seen uptake as a dependency in several software packages, including *SnapATAC2* (18), *ProteinPaint* (19), and *Oxbow* (20)), highlighting its utility.

The *Bigtools* Rust library and CLI executable are available through `crates.io` using Rust’s Cargo package manager:

`$ cargo install bigtools`

The executables are also available on GitHub. *Pybigtools* is available on PyPI:

`$ pip install pybigtools`

*Bigtools* is cross-platform and the executables and Python package are distributed for multiple platforms (including Linux, MacOS, and Windows) and architectures. Documentation for the *Bigtools* library and CLI can be found at https://docs.rs/bigtools and documentation for *Pybigtools* can be found at https://bigtools.readthedocs.io/.

## ACKNOWLEDGEMENTS

We thank Vedat Yilmaz, Peter Kerpedjiev, Trevor Manz, Garrett Ng, and the Weng and Maehr labs for helpful discussions and suggestions. JDH and NA acknowledge funding from the NIH Common Fund 4D Nucleome Program (DK107980).

